# The association between resting state aperiodic activity and Research Domain Criteria Social Processes in young neurotypical adults

**DOI:** 10.1101/2023.09.19.558536

**Authors:** Talitha C. Ford, Aron T. Hill, Nina-Francesca Parrella, Melissa Kirkovski, Peter Donaldson, Peter G. Enticott

**Affiliations:** Cognitive Neuroscience Unit, School of Psychology, Deakin University, Burwood, Victoria, Australia; Centre for Mental Health and Brain Sciences, Swinburne University of Technology, Hawthorn, VIC, Australia; College of Sport, Health and Engineering, Victoria University, Melbourne, Victoria, Australia

**Author notes:** Corresponding author Dr. Talitha Ford, Cognitive Neuroscience Unit, School of Psychology, Deakin University 221 Burwood Hwy, Burwood, Victoria 3125, Australia.

**Keywords:** aperiodic slope, aperiodic exponent, excitation-inhibition, social communication, autism

## Abstract

The aperiodic exponent of the electrophysiological signal has been utilised to demonstrate differences in brain excitation-inhibition in ageing, cognition, and neuro- and psycho-pathology. Furthermore, excitation-inhibition imbalance has been associated with social communication difficulties in clinical and non-clinical cohorts. No work to date, however, has explored the association between aperiodic activity and social communication difficulties. A total of 40 neurotypical young adults aged 18-24 (24 female; age mean = 21.6, SD = 1.9) each underwent 5 minutes of eyes open and eyes closed resting state electroencephalography using a 64-channel HydroCel Geodesic Sensor Net. Participants also completed Research Domain Criteria *social processes* measures: Social Responsiveness Scale, Reading the Mind in the Eyes task, and Penn Emotional Recognition Task. Sex differences in aperiodic exponent and offset were observed, with larger exponent and greater offset observed in males, suggestive of greater inhibitory tone. Whole-brain, as well as left and right social brain, aperiodic exponent were moderately correlated with Social Responsiveness Scale Social Communication Index, but not Reading the Mind in the Eyes or Penn Emotional Recognition tasks. However, the correlations between aperiodic exponent and Social Responsiveness Scale Social Communication Index did not survive multiple comparisons correction, and regression models demonstrated that neither aperiodic exponent or offset were significantly predictive of *social processes* outcomes. These preliminary data provide some evidence that increased aperiodic activity may be associated with social communication difficulties, however, larger samples are needed to validate these initial findings.

## 1. Introduction

Maintaining the equilibrium of excitatory and inhibitory synaptic membrane currents is essential for neural development, information processing, and memory storage (Zhou & Yu, 2018). The brain’s primary inhibitory neurotransmitter, gamma-aminobutyric acid (GABA), is responsible for regulating excitatory glutamate, with disturbance to inhibitory tone leading to hyper-excitation. Likewise, disruption to excitatory glutamatergic processes can result in hyper-excitation depending on the location and nature of the disturbance (Sohal & Rubenstein, 2019). An imbalance in brain excitation/inhibition (E/I) ratios biased toward over-excitation has been associated with several psychopathologies and neurodevelopmental conditions, such as autism spectrum disorder (Bruining et al., 2020; Dickinson et al., 2016; Fatemi, 2008; Gogolla et al., 2009; Nelson & Valakh, 2015; Rubenstein & Merzenich, 2003), schizophrenia (Egerton et al., 2020; Marsman et al., 2013), and epilepsy (Bozzi et al., 2018; Hanada, 2020). Due to the heterogeneity and spectrum nature of clinical conditions such as autism and schizophrenia, it is important to investigate the extent to which brain processes relate to specific symptom domains, rather than within the constraints of diagnostic boundaries. The broader autism phenotype reflects the notion that milder autistic traits are present in varying degrees across the non-clinical population (Austin, 2005; Hurley et al., 2007). Indeed, many of the behavioural and neurophysiological differences observed between autism and control groups appear to hold true in non-clinical high autistic trait groups (Abu-Akel et al., 2017; Ajram et al., 2019; Donaldson et al., 2017; Ford, Nibbs, et al., 2017a; Hill et al., 2025; Kirkovski et al., 2016). Thus, extensive evidence supports the utility of non-clinical populations in investigating the behavioural and neurobiological characteristics of autism (Donaldson et al., 2017; Ford, Nibbs, et al., 2017b; Hill, Van Der Elst, et al., 2022).

Both animal and human studies point firmly toward E/I imbalance as a putative marker of social communication difficulties in clinical and non-clinical populations (Bredewold et al., 2015; Cochran et al., 2015; Ford, Nibbs, et al., 2017b; Han et al., 2014; Yizhar et al., 2011). Human neuroscientific studies support the role of brain E/I in social communication (Ajram et al., 2019; Ford & Crewther, 2016), with increased glutamate and/or reduced GABA levels associated with social communication difficulties in clinical (Brown et al., 2013; Cochran et al., 2015; Gaetz et al., 2014; Hegarty et al., 2018; Pretzsch et al., 2019, 2021; Tebartz van Elst et al., 2014) and non-clinical samples (Ford, Nibbs, et al., 2017b, 2017a; Tebartz van Elst et al., 2014). Furthermore, psychopharmacological interventions that enhance brain inhibition, such as cannabidiol (CBD), riluzole, and R-Baclofen (STX209), have been shown to improve social functioning in children and adults with autism (Ajram et al., 2017; Aran et al., 2021; Cifelli et al., 2020; da Silva Junior et al., 2021; Frye, 2014; Hacohen et al., 2022; Parrella et al., 2023). In mice, optogenetic enhancement of excitation led to impairment in social behaviours, while systematically increasing inhibitory tone thereafter partially rescued these impairments (Yizhar et al., 2011). Similarly, low-frequency repetitive transcranial magnetic stimulation (TMS), which is known to inhibit neural activity, improved autism-like social and behavioural outcomes (Tan et al., 2018). Further, pharmacologically enhancing excitation reduces social behaviours in mice (Bredewold et al., 2015), while enhancing inhibition in mouse models improves social, cognitive, and behavioural outcomes (Han et al., 2014; Stoppel et al., 2018). Taken together, these studies point firmly toward an E/I imbalance contributing to social communication difficulties across clinical lines.

Brain E/I imbalance can disrupt social communication abilities in several ways (Parrella et al., 2024), particularly in social brain regions such as the superior temporal sulcus (STS), temporo-parietal junction (TJP), and medial prefrontal cortex (mPFC) (Blakemore, 2008; Frith & Frith, 2007). E/I imbalance in social brain regions can affect connectivity between brain networks that are responsible for processing social cues (Gao & Penzes, 2015) and sensory sensitivities that heighten anxiety in social situations (Balasco et al., 2020; A. M. Gonçalves & Monteiro, 2023).

Electroencephalography (EEG) has long been utilised as an accessible and affordable tool to probe the relationship between brain and behaviour, particularly in relation to psychiatric and neurodevelopmental conditions (Milovanovic & Grujicic, 2021). Putative markers of excitation and/or inhibition include mismatch negativity (Greenwood et al., 2018; Rowland et al., 2016; Sehatpour et al., 2022), functional E/I ratio (Bruining et al., 2020), and oscillatory activity in delta, theta, and gamma bands (Barttfeld et al., 2011; Coben et al., 2008; Grent-’t-Jong et al., 2018; Lisman, 2012; Uhlhaas & Singer, 2010). Recent developments in resting state and task-related EEG data analysis have revealed that the broadband aperiodic signal, in contrast to periodic activity within a narrowband frequency range (i.e., neuronal oscillations), might be a more accurate reflection of E/I balance (Donoghue et al., 2020; Gao et al., 2017; Waschke et al., 2021). Aperiodic activity is the background, scale-free activity that is represented on a 1/f-like power distribution, where spectral power decreases as frequency increases (Donoghue et al., 2020; Hill, Clark, et al., 2022).

The aperiodic signal, which is comprised of both the exponent (spectral slope) and offset (broadband shift in power across frequencies), is thought to reflect the integration of underlying synaptic currents (Donoghue et al., 2021). These currents have a double-exponential shape in the time domain, which produces the 1/f-like spectrum (Donoghue et al., 2020; Gao et al., 2017). Excitatory currents have a faster time constant and produce constant power at lower frequencies before rapid decay, while power decays more slowly as a function of frequency for inhibitory currents. Thus, a smaller exponent (flatter 1/f-like slope) is thought to reflect more excitation relative to inhibition, while a larger exponent (steeper slope) reflects the converse (Donoghue et al., 2020; Gao et al., 2017). The aperiodic offset (i.e., the y-intercept of the spectrum) is thought to reflect neuronal population spiking (Manning et al., 2009; Miller et al., 2014), which is regulated by GABAergic interneurons (Tremblay et al., 2016). Although the neural underpinnings of the offset are not well understood, offset and exponent are highly interrelated and thus correlated, suggesting that the offset, at least in part, is reflective of excitatory and/or inhibitory processes. Aperiodic exponent and offset have been associated with age (Cellier et al., 2021; Donoghue et al., 2020; Hill, Clark, et al., 2022; McSweeney et al., 2021), sex (McSweeney et al., 2021), and several cognitive processes such as working memory (Donoghue et al., 2020), visual attention (Waschke et al., 2021), and meta control (Zhang et al., 2023). It is important to note, however, that the exact physiological mechanisms that underlie changes in aperiodic activity are likely complex and remain subject to ongoing investigation. Therefore, interpretation of the functional relevance of aperiodic exponent and offset should be done with caution at this stage. For recent reviews of aperiodic activity within clinical and neurodevelopmental contexts see (Pani et al., 2022; Stanyard et al., 2024).

Given its utility for examining E/I balance across the cortex, and relative affordability and simple acquisition, the aperiodic exponent has been utilised to demonstrate E/I differences in autism (Arutiunian et al., 2024), schizophrenia (Molina et al., 2020), ADHD (Ostlund et al., 2021; Robertson et al., 2019), and Fragile X Syndrome (Wilkinson & Nelson, 2021). To our knowledge, three studies to date have investigated aperiodic exponent in relation to social communication: social-emotional development in toddlers born preterm, with increased aperiodic exponent (increased inhibition) associated with more social emotional problems (Shuffrey et al., 2022); toddlers with neurofibromatosis, with increased exponent associated with higher autistic traits for those with poor executive function (Carter Leno et al., 2022); and typically developing children with autistic traits associated with a larger exponent, when EEG was recorded during a facial emotion processing task (Hill et al., 2025). No work to date has applied aperiodic activity analyses to explore differences in E/I and social communication in a non-clinical adult population.

The Research Domain Criteria (RDoC) highlight the limitations of diagnosis-specific research, and provides a rich framework for understanding psychiatric conditions in the context of varying degrees of symptomatology with neurobiological and/or genetic origins (Insel, 2010). Within the RDoC, the *social processes* domain includes social communication, affiliation and attachment with others, and perception and understanding of the self and others (Cuthbert & Insel, 2013). Self-report measures of *social processes* recommended by RDoC include the Social Responsiveness Scale (SRS; Constantino & Gruber, 2005), while objective measures include the Penn Emotion Recognition (ER-40; Gur et al., 2002), and Reading the Mind in the Eyes (RME; Baron-Cohen et al., 2001) tasks. These measures have been utilised to assess social functioning in both clinical and non-clinical populations. Understanding the neurobiological mechanisms of *social processes*, at both clinical and non-clinical levels, is essential for broader global efforts to identify effective therapeutics to support those with social processing difficulties.

To that end, this study explores the extent to which aperiodic activity predicts RDoC *social processes* outcomes in young non-clinical adults. In line with the brain E/I hypothesis in autism, we hypothesised that a reduced aperiodic exponent (flatter slope), indicating reduced global inhibitory tone (E>I), would predict more *social processes* difficulties. Given there has been no literature published to date on aperiodic activity and social difficulties, we also explored this hypothesis specifically in right and left temporo-parietal “social brain” regions, which are known to be involved in social information processing (Blakemore, 2008; Frith & Frith, 2007).

## 2. Methods and Materials

### 2.1. Participants

A total of 40 young adults (24 female, 16 male) aged 18-24 participated in this study (mean age = 21.6 years, *SD* = 1.9). Participants had an IQ > 90 (range 91-148, mean = 114.6, *SD* = 10.15), measured using the Wechsler Abbreviated Scales of Intelligence (version 2; WASI-III)(Wechsler, 2011), reflecting “average” (90-109) through to “superior” (>130) intelligence within the sample. Education levels included 7 participants with secondary college, 29 with a tertiary degree (undergraduate, diploma, vocational training), and 4 with postgraduate degree (PhD, Masters). Represented ethnicities were: 30 European, 5 East Asian, 4 South Asian, 1 Middle Eastern. Participants were excluded if they self-reported: a history of developmental, psychiatric genetic, or neurological conditions; a first-degree relative with a history of autism spectrum disorder, schizophrenia, bipolar disorder, or other psychiatric condition (not including depression and anxiety); a substance use disorder; regular smoking (> 1 cigarette/day); or taking any psychoactive medications. Participants were asked to abstain from caffeine and nicotine for 12 hours, and recreational drugs for 1 week, prior to testing; confirmation of compliance was made verbally on the day of testing. Participants provided written informed consent prior to any data collection for this study and were reimbursed $30AUD for their time participating in the study. Ethics approval was granted by Deakin University’s Human Research Ethics Committee (2019-208) in accordance with the Declaration of Helsinki.

### 2.2. Psychometric and Behavioural Measures

The below measures were administered prior to EEG data collection.

*Social Responsiveness Scale Version-2* (SRS; Constantino & Gruber, 2012): The SRS is a 65-item measure of symptom severity across five autism symptom domains: Social Awareness (8 items), Social Motivation (11 items), Social Cognition (12 items), Social Communication (22 items), and Repetitive Behaviours/Restricted Interests (12 items). Responses are measured on a 4-point scale from 1 (*not true*) to 4 (*almost always true*), and t-scores are calculated from raw scores to provide an indication of severity of autistic symptoms (“normal”: <60, “mild”: 60-65, “moderate”: 66-75, “severe”: >75). The psychometric properties of the adult self-report measure of the SRS were assessed by Ingersoll et al. (2011), with good reliabilities across subscales reported in an adult sample (SRS total α = .95, Social Awareness α = .64, Social Cognition α = .78, Social Communication α = .88, Social Motivation α = .81, and Repetitive Behaviours/Restricted Interests α = .81). Within the RDoC framework, the SRS is a measure of Social Communication – reception and production of non-facial communication. Given our focus on social communication, the SRS Social Communication Index (SRS SCI) was utilised, which is a composite score of subscales: Social Awareness, Social Motivation, Social Cognition, and Social Communication (Constantino & Gruber, 2005, 2012).

*Reading the Mind in the Eyes Task* (RME; Baron-Cohen et al., 2001): The RME is a 36-item measure of perception and understanding of conspecific mental states through the assessment an actor’s eyes. The computerised task presents consecutive photos of actor’s eyes in the centre of the screen, and four words (one in each corner of the screen) that might describe the emotional state of the eye. Participants are to select the word that best describes the actor’s emotion via mouse click. Accuracy on the RME is scored as the proportion of correct responses (from 0 to 1). The RME is widely used, and has been shown to exhibit acceptable internal consistency (α = .73)(Kittel et al., 2022). Because participants are provided a dictionary of the emotional labels presented, reaction time data are not analysed. Within the RDoC framework, the RME is a measure of Perception and Understanding of Others – understanding mental states.

*Penn Emotion Recognition Task* (ER-40; Gur et al., 2002): The ER-40 consists of 40 colour photographs of actors expressing one of four emotions (happy, sad, anger, or fear) or neutral expressions. Stimuli are balanced for actors’ gender, age, and ethnicity, and for each emotion category. Four high-intensity and four low-intensity expressions are included for each emotion. Participants view one photograph at a time and choose the correct emotion from a list to the right-hand side of the face via mouse click. The ER-40 has shown adequate test-retest reliability (*r* = .753) and internal consistency (α = .645) in non-clinical adults (Pinkham et al., 2016).

A composite measure of accuracy and reaction time, the linear integrated speed-accuracy score (LISAS), is calculated according to the following formula: LISAS = RT + (S_RT_ / S_PE_) x PE, where RT is participant mean reaction time, PE is participant mean proportion of errors, S_RT_ is the standard deviation of the reaction time, and S_PE_ is the standard deviation of the proportion or errors (Vandierendonck, 2017). ER-40 LISAS will be referred to as “ER-40 performance” hereafter, with higher scores indicating better performance. Within the RDoC framework, the ER-40 task is a measure of Social Communication – reception of facial communication.

### 2.3. Electroencephalography data acquisition

EEG data were recorded using a 64-channel HydroCel Geodesic Sensor Net (Electrical Geodesics, Inc, USA) containing silver/silver-chloride (Ag/AgCl) electrodes surrounded by electrolyte-wetted sponges. Electrical Geodesics Net Station EEG software (version 5.0) was used to acquire the data via a Net Amps 400 amplifier using a sampling rate of 1 KHz. Data were referenced to electrode Cz and grounded to the common of the isolated amp circuit’s power supply. Electrode impedances were checked to ensure they were < 50 KOhms prior to data collection. Participants were seated in a quiet, dimly lit room and asked to sit still with their eyes open while looking at a fixation cross for five minutes, and then eyes closed for five minutes, while resting state data were collected. Electrode impedances were checked between eyes open and eyes closed blocks to ensure they remained < 50 KOhms (considered low impedance for this system). One participant withdrew from the study and did not undergo EEG (remaining *n* = 39).

### 2.4. EEG pre-processing

EEG data were pre-processed in MATLAB (R2021a; The Mathworks, Massachusetts, USA) incorporating the EEGLAB toolbox (v2023.0; Delorme & Makeig, 2004) and custom scripts. The Reduction of Electroencephalographic Artifacts (RELAX; v1.1.2) software (Bailey, Biabani, et al., 2023; Bailey, Hill, et al., 2023) was used to clean each EEG dataset. The fully automated RELAX pipeline uses empirical approaches to identify and reduce non-neural artifacts, and includes the use of both multi-channel Wiener filters and wavelet enhanced independent component analysis (ICA). As part of the RELAX pipeline, the data were bandpass filtered between 0.5 and 80 Hz (fourth-order zero-phase Butterworth filter). A notch filter (47-53 Hz) was then applied to remove any line noise. Any bad channels were identified and removed using a multi-step process incorporating the default parameters of the ‘findNoisyChannels’ function (PREP pipeline; Bigdely-Shamlo et al., 2015). Multi-channel Wiener filtering (Castellanos & Makarov, 2006) was used to initially clean blinks, muscle activity, and horizontal eye movement, followed by robust average re-referencing (Bigdely-Shamlo et al., 2015), and wavelet-enhanced ICA (Castellanos & Makarov, 2006), with components for cleaning identified using the automated independent component (IC) classifier IClabel (Pion-Tonachini et al., 2019). Any rejected electrodes were then interpolated back into the data using spherical interpolation and all pre-processed data files were visually inspected prior to further analysis (Eyes closed data: mean rejected channels = 4.89, *SD* = 3.52; Eyes open data: mean rejected channels = 7.75, *SD* = 3.72).

### 2.5. Electroencephalography data analysis

All pre-processed resting state data were visually inspected for quality assurance prior to inclusion in statistical analyses. Two participants were excluded from the study due to excessive electrical line noise for both eyes open and eyes closed conditions (remaining *n* = 37) that persisted after cleaning, while an additional two participants’ eyes open data were excluded due to excessive line noise (eyes open *n* = 35). The average length of continuous data (in seconds) after cleaning for the eyes open condition was 299.27 seconds (*SD* = 25.13), and for the eyes-closed condition was 304.70 seconds (*SD* = 32.29). The power spectral density of the EEG signal was calculated using Welch’s method (2-second Hamming window, 50% overlap) for all electrodes across all participants for both the eyes-open and eyes-closed datasets. The specparam (formally FOOOF) toolbox (version 1.0.0; https://fooof-tools.github.io/fooof/; Donoghue et al., 2020) was then used to parameterise the spectral data through separation of the periodic and aperiodic components of the signal. The aperiodic exponent and offset were extracted across 1 to 40 Hz using the ‘fixed’ aperiodic mode. Spectral parameterization settings for the algorithm were based on recommendations on the specparam toolbox website (https://fooof-tools.github.io/fooof/) and past literature (Ostlund et al., 2022): peak width limits = [1, 12], maximum number of peaks = 8, peak threshold = 2, minimum peak height = .05.

The performance of the FOOOF algorithm for fitting the data was assessed using two “goodness of fit” measures: R^2^, explained variance; and mean absolute error (MAE), total error of the model fit (Donoghue et al., 2020; Ostlund et al., 2021). Model fits for both eyes open and eyes closed data were good (eyes open: R^2^ = .99, MAE = .047; eyes closed: R^2^ = .99, MAE = .054. Complete fit statistics are presented in the Supplementary Table 1). There was no difference in mean R^2^ between eyes open and eyes closed conditions (*t*(64.13) = -.30, *p* = .765), while MAE for eyes open was significantly lower compared to eyes closed (*t*(69.44) = 2.19, *p* = .032). Aperiodic exponent (slope) was significantly larger for the eyes closed condition, compared to the eyes open condition (*t*(69.65) = 2.82, *p* = .006). Similarly, the offset (intercept) was larger for eyes closed than eyes open condition (*t*(69.97) = 3.0, *p* = .004). There were strong, positive correlations between exponent and offset values for eyes open (*rho* = .73, *p* < .001) and eyes closed (*rho* = .80, *p* < .001) recordings, and between eyes open and closed exponent (*rho* = .73, *p* < .001) and eyes open and closed offset (*rho* = .79, *p* < .001).

To calculate aperiodic exponent and offset in brain regions associated with social information processing, so called “social brain” (Blakemore, 2008; Moessnang et al., 2016), electrodes over left and right temporo-parietal regions were averaged separately (left: P5 CP5 TP9 TP7 C5 T7 T9; right: P6 CP6 TP10 TP8 C6 T8 T10; see Figure 1). These electrodes cover brain regions including the superior, middle, and inferior temporal gyrus, supramarginal gyrus, and angular gyrus, all known to be involved in social information processing (Blakemore, 2008; Jung et al., 2019; Moessnang et al., 2016).

**Figure 1:**
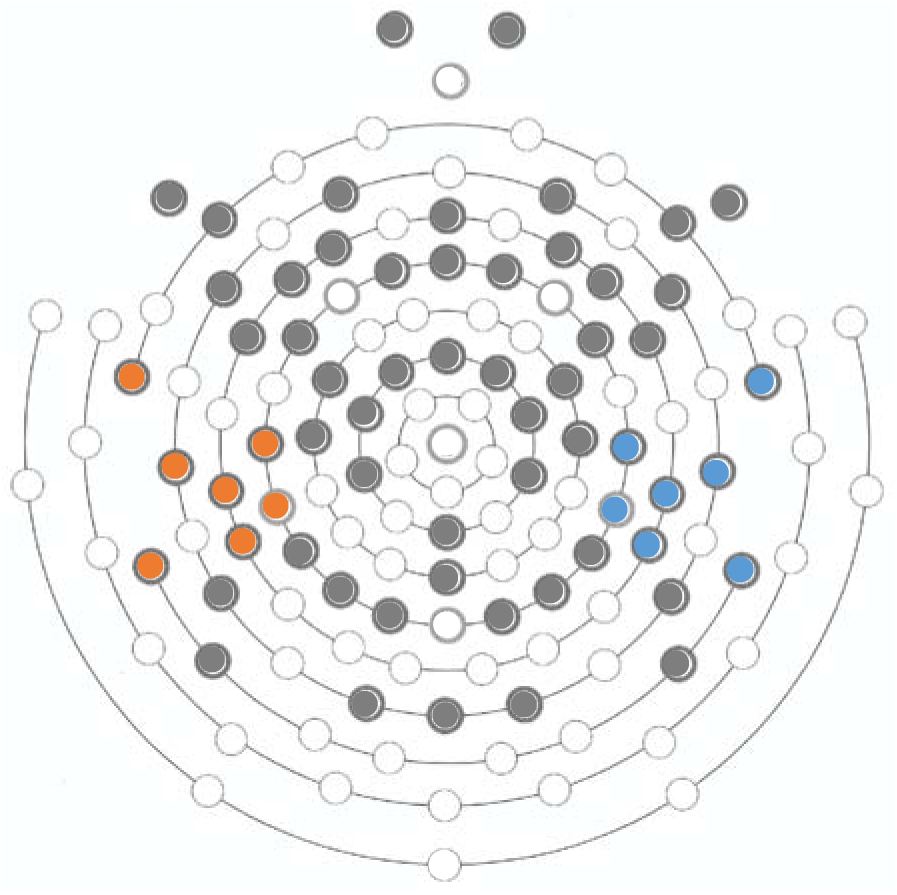
Electroencephalograph electrode placement. 64-channel Geodesic Sensor Net electrode placement (dark circles), with the cluster of electrodes selected to represent left (blue) and right (orange) social brain highlighted.

### 2.6. Statistical analyses

Statistical analyses were conducted in R (version 4.1.0). Given this is the first study to examine aperiodic activity in relation to social information processing, *a priori* power analyses were not conducted. However, previous studies have reported significant associations between brain E/I and social communication with sample sizes of 29 to 37 non-clinical adults (Ford, Nibbs, et al., 2017a; Ford, Woods, et al., 2017; Tebartz van Elst et al., 2014). Assumptions for all statistical analyses were assessed using the *olsrr* and *MVN* packages in R (Aravind Hebbali, 2020; Korkmaz et al., 2014), including normality, linearity, homoscedasticity, and multicollinearity. All assumptions were met, except for a significant positive skew in RME and negative skew in ER-40 LISAS scores, which were subsequently normalised using the Yeo-Johnson transform in R (VGAM package; Yeo & Johnson, 2000).

Initial bivariate Pearson’s r correlation analyses were conducted to investigate the association between *social processes* outcomes (SRS SCI score, RME accuracy, and ER-40 performance) and whole-brain aperiodic exponent and offset with eyes open and eyes closed. Additional exploratory correlations analyses were conducted to investigate the association between *social processes* outcomes and left and right social brain exponent and offset. The significance level was set at *p* = .05, and false discovery rate (FDR) was corrected for using the Benjamin and Hochberg method in R (Benjamini & Hochberg, 1995; stats package; R Core Team, 2023).

Then, multivariate linear regressions (MLRs) were conducted to assess the extent to which aperiodic exponent and offset with eyes open and eyes closed predicted *social processes* outcomes. Due to the high correlation between EO and EC exponent and offset, four separate MLR models were defined (i.e., model 1 predictor: eyes open exponent; model 2 predictor: eyes closed exponent; model 3 predictor: eyes open offset; model 4 predictor: eyes closed offset). There were significant sex effects for exponent and offset in both conditions, therefore, sex was entered as a covariate for each model (see Supplementary Material for full model designs). Additional exploratory MLR models were defined to investigate the extent to which aperiodic activity in left and right social brain predicted *social processes* outcomes; again, sex was entered as a covariate (see Supplementary Material for full model designs). The significance level for each model was set at *p* = .05, and FDR was corrected for using the Benjamini and Hochberg method in R (Benjamini & Hochberg, 1995; stats package; R Core Team, 2023).

## 3. Results

### 3.1. Descriptive statistics

Descriptive statistics for final participant sample characteristics (*n =* 37), *social processes* outcomes, and aperiodic activity metrics are presented in Table 1. There were no significant age or biological sex effects on *social processes* measures (*p*s > .05), and no significant correlations between age and aperiodic activity outcomes (*p*s > .05). However, males compared to females had a significantly larger exponent (*t*(24.20) = -2.56, *p* = .017, *p_FDR_* = .033) and higher offset (*t*(29.20) = -2.08, *p* = .047, *p_FDR_* = .047) during eyes open, as well as a larger exponent (*t*(33.76) = -2.83, *p* = .008, *p_FDR_* = .032) and higher offset during eyes closed (*t*(34.90) = -2.34, *p* = .025, *p_FDR_* = .033; Figure 2).

**Figure 2:**
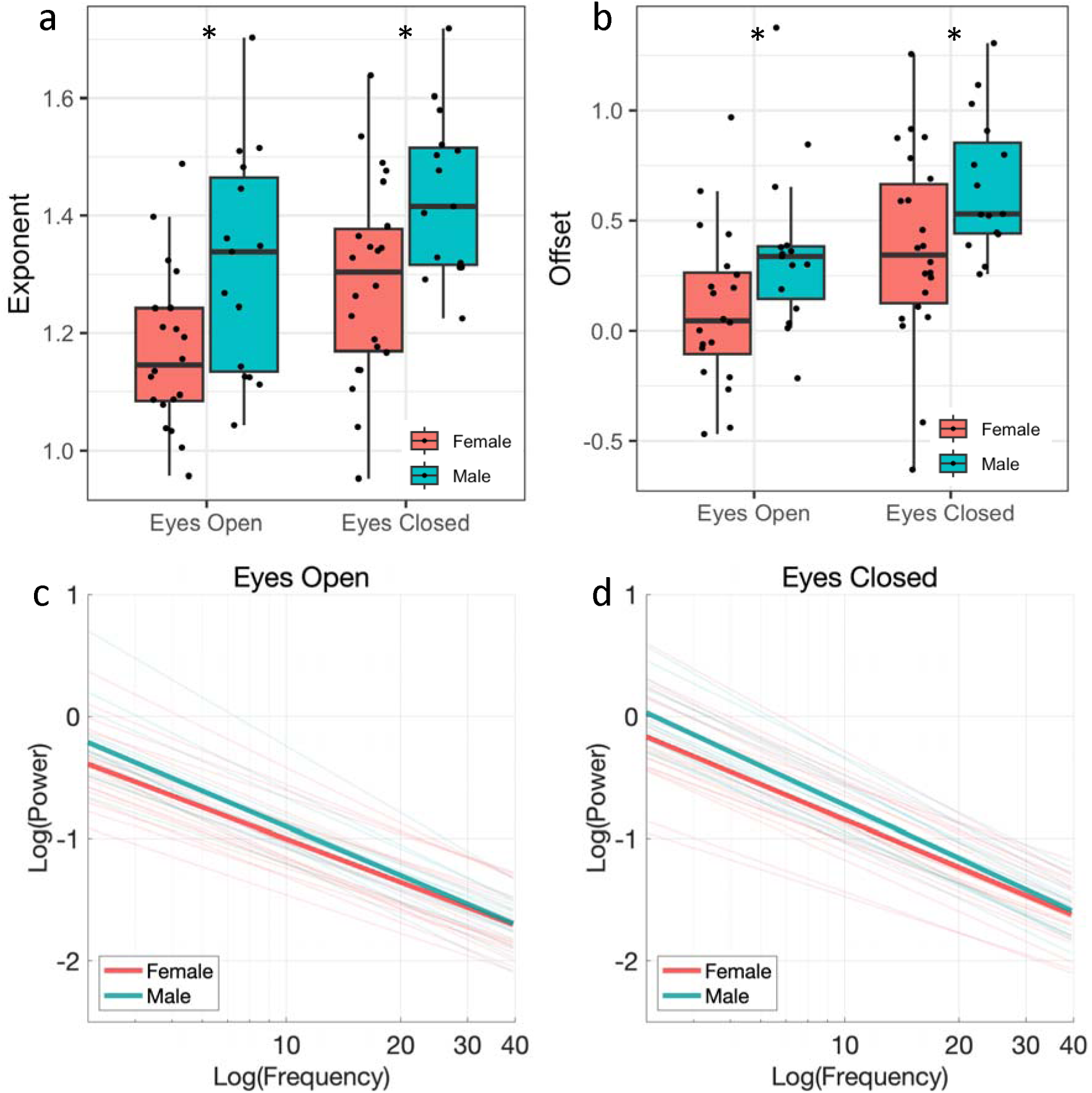
Sex differences in aperiodic activity. Sex differences in exponent (a) and offset (b) for eyes open and eyes closed conditions, and power vs frequency plots by sex for eyes open (c) and eyes closed (d) conditions showing the aperiodic component of the neural signal. Thin lines represent the aperiodic signal for individual participants, thick lines represent the average across all participants. \**p_FDR_*<.05

**Table 1:** Descriptive statistics of psychometric, behavioural, and aperiodic activity measures.

|  | Total<br>(*n* = 37) |  | Female<br>(*n* = 22) | Male<br>(*n* = 15) |
| --- | --- | --- | --- | --- |
|  | Mean (SD) | Min-Max | Mean (SD) | Mean (SD) |
| Age | 22 (1.9) | 18 – 25 | 21 (1.9) | 22 (1.9) |
| WASI IQ | 115 (9.8) | 91 – 148 | 114 (8.2) | 117 (12) |
| *Social Processes* |  |  |  |  |
| SRS total t-score | 52 (6.8) | 42 – 69 | 51 (5.8) | 54 (8.4) |
| SRS SCI t-score | 53 (6.9) | 42 – 71 | 52 (5.6) | 54 (8.6) |
| SRS RRB t-score | 53 (8.4) | 40 – 73 | 51 (7.2) | 55 (10) |
| RME accuracy | .81 (.08) | .64 – .94 | .83 (.08) | .78 (.08) |
| ER-40 RT (sec) | 2.9 (.69) | 2.0 – 4.9 | 2.8 (.48) | 3.0 (.94) |
| ER-40 accuracy | .89 (.05) | .78 – .97 | .90 (.04) | .86 (.05) |
| ER-40 performance | 3.3 (.97) | 2.2 – 7.0 | 3.1 (.65) | 3.5 (1.3) |
| *Aperiodic activity* |  |  |  |  |
| *Whole brain* |  |  |  |  |
| EO exponent (a.u.) | 1.2 (.17) | .96 – 1.7 | 1.2 (.14) | 1.3 (.19) |
| EC exponent (a.u.) | 1.3 (.17) | .95 – 1.7 | 1.3 (.17) | 1.4 (.14) |
| EO offset ($\mu V^2$) | .21 (.38) | -.47 – 1.4 | .10 (.36) | .36 (.38) |
| EC offset ($\mu V^2$) | .49 (.41) | -.63 – 1.3 | .37 (.44) | .66 (.31) |
| *Left social brain* |  |  |  |  |
| EO exponent (a.u.) | 1.2 (.19) | .96 – 1.7 | 1.2 (.15) | 1.3 (.21) |
| EC exponent (a.u.) | 1.3 (.20) | .86 – 1.7 | 1.3 (.20) | 1.4 (.15) |
| EO offset ($\mu V^2$) | .16 (.41) | -.50 – 1.5 | .014 (.34) | .35 (.43) |
| EC offset ($\mu V^2$) | .42 (.45) | -.67 – 1.3 | .25 (.45) | .66 (.35) |
| *Right social brain* |  |  |  |  |
| EO exponent (a.u.) | 1.2 (.20) | .67 – 1.7 | 1.2 (.18) | 1.3 (.19) |
| EC exponent (a.u.) | 1.3 (.20) | .77 – 1.6 | 1.2 (.21) | 1.4 (.13) |
| EO offset ($\mu V^2$) | .18 (.42) | -.84 – 1.3 | .026 (.37) | .38 (.41) |
| EC offset ($\mu V^2$) | .43 (.48) | -1.0 – 1.2 | .26 (.50) | .67 (.34) |

Notes: SD = standard deviation, WASI IQ = Weschler Abbreviated Scale of Intelligence Intelligence Quotient, SRS = Social Responsiveness Scale, SCI = Social Communication Index, RRB = Restricted Repetitive Behaviours, RME = Reading the Mind in the Eyes, ER-40 = Penn Emotion Recognition Task, RT = reaction time, performance = linear integrated speed- accuracy score (LISAS), EO = eyes open, EC = eyes closed.

### 3.2. Social processes outcomes

Pearson’s r correlations were conducted to test the relationship between the three *social processes* measures. There were no significant relationships between SRS SCI scores and RME accuracy or ER-40 performance (*p_FDR_* > .05). The relationship between ER-40 performance and RME accuracy was significant, with ER-40 performance increasing with RME accuracy as would be expected (*r* = .434, *p* = .018, *p_FDR_* = .025).

### 3.3. Correlations between aperiodic activity and *social processes* outcomes

Pearson’s r correlations between *social processes* measures and whole-brain and social brain aperiodic activity are presented in Table 2. There were significant moderate correlations between SRS SCI score and whole brain exponent, left social brain exponent and offset, and right social brain exponent in the eyes open condition only (uncorrected p < .05), with aperiodic activity increasing with more social communication difficulties. However, these correlations did not survive FDR correction (p_FDR_ > .05). There were no significant correlations between RME accuracy and ER-40 performance and aperiodic activity across whole-brain or social brain regions (uncorrected p > .05). Scatterplots for these associations are presented in Figure 3 and Figure 4.

**Figure 3:**
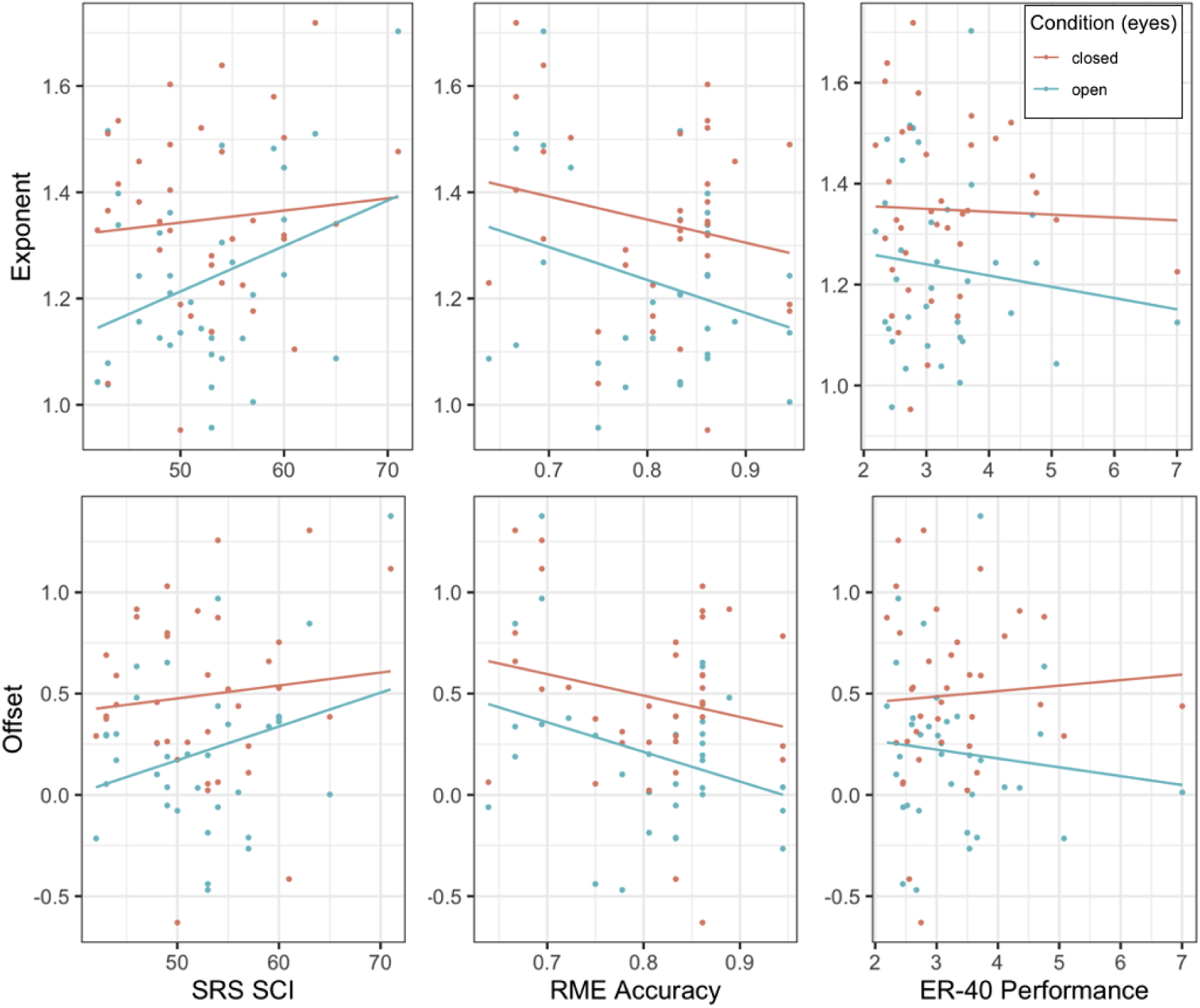
Scatter plots of aperiodic activity and Social Processes outcomes. Scatterplots of depicting exponent (top row; model 1 & 2) and offset (bottom row; model 3 & 4) versus Social Responsiveness Scale Social Communication Index score (SRS SCI; left panel), Reading the Mind in the Eyes (RME) accuracy (middle panel), and Penn Emotion Recognition (ER-40) performance (right panel) in eyes open (green) and eyes closed (red) conditions.

**Figure 4:**
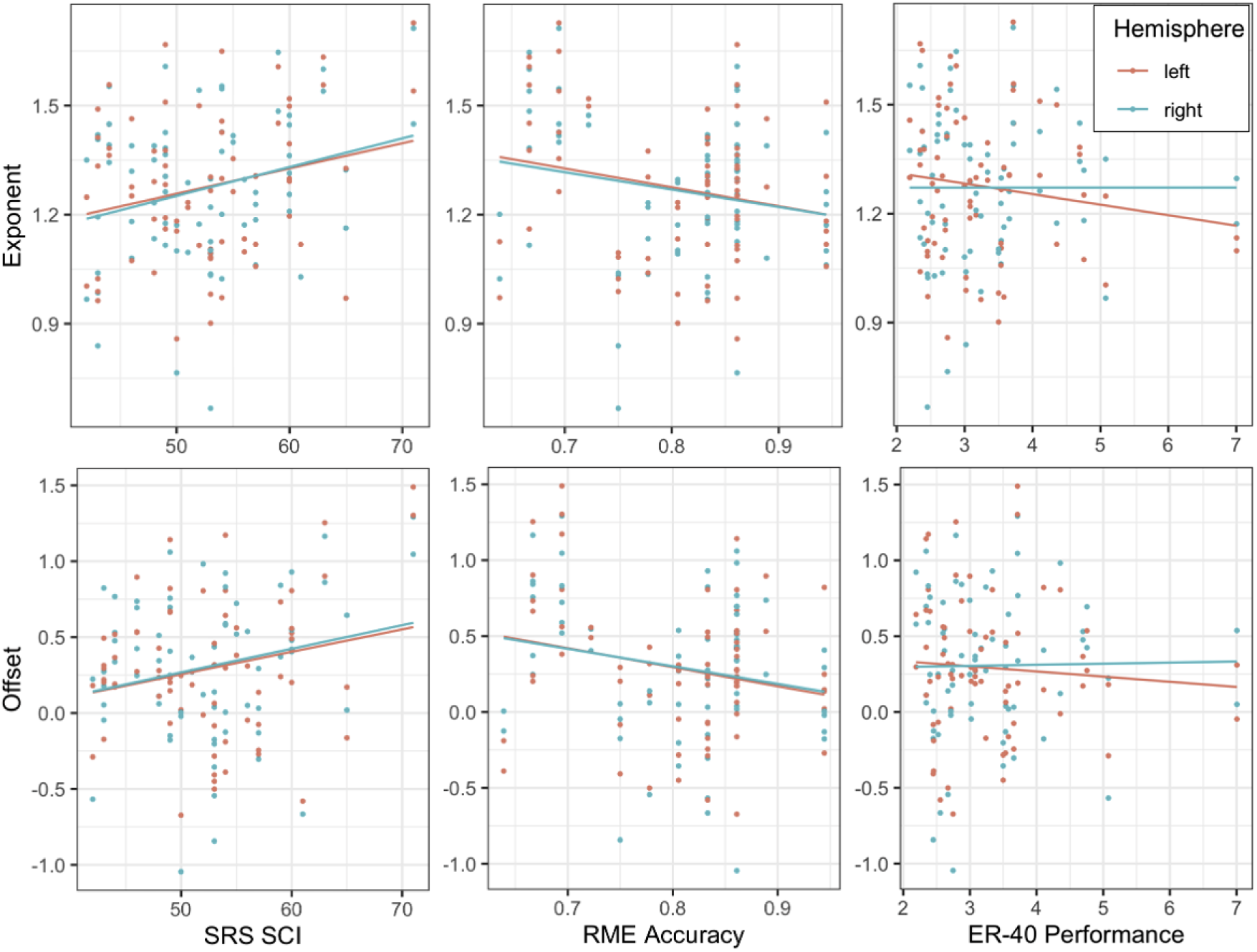
Scatter plots of aperiodic activity by hemisphere and Social Processes outcomes. Scatter plots of exponent (top row; model 1 & 3) and offset (bottom; model 2 & 4) versus Social Responsiveness Scale Social Communication Index score (SRS SCI; left panel), Reading the Mind in the Eyes (RME) accuracy (middle panel), and Penn Emotion Recognition (ER-40) performance (right panel) in left (red) and right (green) social brain regions.

**Table 2:** Pearson’s r correlations between RDoC *social processes* measures and aperiodic activity.

|  | SRS SCI score | RME accuracy | ER-40 performance |
| --- | --- | --- | --- |
| *Whole brain* |  |  |  |
| EO Exponent (a.u.) | .34<sup>a</sup> | -.29 | -.09 |
| EC Exponent (a.u.) | .09 | -.20 | -.04 |
| EO Offset ($\mu V^2$) | .30 | -.32 | -.11 |
| EC Offset ($\mu V^2$) | .10 | -.21 | .06 |
| *Left social brain* |  |  |  |
| EO Exponent (a.u.) | .35<sup>a</sup> | -.23 | -.12 |
| EC Exponent (a.u.) | .14 | -.21 | -.12 |
| EO Offset ($\mu V^2$) | .34<sup>a</sup> | -.27 | -.14 |
| EC Offset ($\mu V^2$) | .14 | -.21 | -.01 |
| *Right social brain* |  |  |  |
| EO Exponent (a.u.) | .40<sup>a</sup> | -.22 | -.04 |
| EC Exponent (a.u.) | .14 | -.17 | .04 |
| EO Offset ($\mu V^2$) | .33 | -.24 | -.10 |
| EC Offset ($\mu V^2$) | .15 | -.19 | .13 |

Notes: SRS = Social Responsiveness Scale, SCI = Social Communication Index, RME = Reading the Mind in the Eyes, ER-40 = Penn Emotion Recognition Task, EO = eyes open, EC = eyes closed. <sup>a</sup>correlations significant at $p < .05$ prior to correction.

### 3.4. Whole-brain aperiodic activity and *social processes* outcomes

Multivariate linear regression models were designed to test the extent to which aperiodic exponent and offset predicted *social processes* outcomes of SRS SCI scores, RME accuracy, and ER-40 performance. Four models were defined with exponent and offset for eyes open and eyes closed entered as separate predictors. None of the models were significant (uncorrected p > .05) suggesting that neither exponent or offset predicted *social processes* outcomes in either eyes open or eyes closed conditions (see Table 3 for model statistics). Figure 3 illustrates the relationships between each of predictor and outcome measures, and suggests a visual trend toward increased exponent (i.e., increased inhibitory tone) and offset with higher SRS SCI scores, and poorer RME and ER-40 performance, particularly in the eyes open condition.

**Table 3:** Multivariate linear regression model results for whole-brain aperiodic activity and RDoC *social processes* measures.

| | $R^2$ | Adj. $R^2$ | $F$ (df) | $p_{uncorr}$ | $p_{FDR}$ | Cohens $f^2$ |
| --- | --- | --- | --- | --- | --- | --- |
| **Exponent** |  |  |  |  |  |  |
| *Model 1: Eyes open* |  |  |  |  |  |  |
| SRS SCI | .119 | .063 | 2.10 (2,31) | .139 | .334 | .119 |
| RME accuracy | .131 | .075 | 2.43 (2,31) | .113 | .334 | .151 |
| ER-40 performance | .008 | -.056 | .23 (2,31) | .881 | .956 | .008 |
| *Model 2: Eyes closed* |  |  |  |  |  |  |
| SRS SCI | .038 | -.020 | .07 (2,33) | .530 | .795 | .039 |
| RME accuracy | .115 | .061 | 2.15 (2,33) | .133 | .334 | .130 |
| ER-40 performance | .003 | -.058 | .05 (2,33) | .956 | .956 | .003 |
| **Offset** |  |  |  |  |  |  |
| *Model 3: Eyes open* |  |  |  |  |  |  |
| SRS SCI | .103 | .045 | 1.79 (2,31) | .185 | .370 | .115 |
| RME accuracy | .148 | .093 | 2.70 (2,31) | .083 | .334 | .174 |
| ER-40 performance | .008 | -.056 | .12 (2,31) | .887 | .956 | .008 |
| *Model 4: Eyes closed* |  |  |  |  |  |  |
| SRS SCI | .039 | -.020 | .66 (2,33) | .522 | .795 | .040 |
| RME accuracy | .119 | .065 | 2.22 (2,33) | .124 | .334 | .135 |
| ER-40 performance | .012 | -.050 | .21 (2,33) | .815 | .956 | .013 |

Notes: Adj = Adjusted, df = degrees of freedom, uncorr = uncorrected, FDR = false discovery rate, SRS = Social Responsiveness Scale, SCI = Social Communication Index, RME = Reading the Mind in the Eyes, ER-40 = Penn Emotion Recognition Task, RDoC = Research Domain Criteria.

### 3.5. Social brain aperiodic activity and social processes outcomes

Exploratory analyses were conducted to investigate whether aperiodic activity in temporal-parietal regions known to be involved in social information processing (Blakemore, 2008; Jung et al., 2019; Moessnang et al., 2016) might predict *social processes* outcomes (SRS SCI, RME accuracy, ER-40 performance). Given the lack of effects of the eyes closed condition in the whole-brain analyses, only eyes open data were analysed here. Therefore, measures of exponent and offset for eyes open were averaged over temporo-parietal clusters in left and right hemisphere, and four models were defined for left and right exponent and offset separately (i.e., left exponent, left offset, right exponent, right offset) (see Supplementary Material for models). Figure 4 illustrates the relationships between *social processes* outcomes and predictors. None of the overall models for SRS SCI, ER-40 performance, and RME accuracy for both exponent and offset predictors were significant (uncorrected *p*s > .05, see Table 4 for model statistics). Despite the non-significant finding, there was a trend toward increased right social brain exponent (i.e., increased inhibitory tone) with higher SRS SCI scores (p = .068), with a moderate effect size (f = .189). When visualising the data in Figure 4, exponent and offset appeared to increase with higher SRS SCI scores and poorer RME accuracy over both social brain regions.

**Table 4:** Multivariate linear regression model results for left and right social brain aperiodic activity and RDoC social processes measures.

| | $R^2$ | Adj. $R^2$ | $F$ (df) | $p_{uncorr}$ | $p_{FDR}$ | Cohen's $f^2$ |
| --- | --- | --- | --- | --- | --- | --- |
| **Exponent** |  |  |  |  |  |  |
| Model 1: Left |  |  |  |  |  |  |
| SRS SCI | .129 | .072 | 2.29 (2,31) | .118 | .251 | .148 |
| RME accuracy | .114 | .056 | 1.99 (2,31) | .154 | .251 | .128 |
| ER-40 performance | .013 | -.050 | .208 (2,31) | .813 | .976 | .013 |
| Model 2: Right |  |  |  |  |  |  |
| SRS SCI | .159 | .105 | 2.94 (2,31) | .068 | .251 | .189 |
| RME accuracy | .109 | .052 | 1.90 (2,31) | .167 | .251 | .123 |
| ER-40 performance | .000 | -.064 | .00 (2,31) | .998 | .998 | .000 |
| **Offset** |  |  |  |  |  |  |
| Model 3: Left |  |  |  |  |  |  |
| SRS SCI | .127 | .064 | 2.13 (2,31) | .136 | .251 | .137 |
| RME accuracy | .124 | .067 | 2.20 (2,31) | .128 | .251 | .142 |
| ER-40 performance | .020 | -.043 | .32 (2,31) | .729 | .972 | .021 |
| Model 4: Right |  |  |  |  |  |  |
| SRS SCI | .114 | .057 | 2.00 (2,31) | .152 | .251 | .129 |
| RME accuracy | .115 | .057 | 2.01 (2,31) | .152 | .251 | .129 |
| ER-40 performance | .003 | -.061 | .054 (2,31) | .948 | .998 | .003 |

Notes: Adj = Adjusted, df = degrees of freedom, uncorr = uncorrected, FDR = false discovery rate, SRS = Social Responsiveness Scale, SCI = Social Communication Index, RME = Reading the Mind in the Eyes, ER-40 = Penn Emotion Recognition Task, RDoC = Research Domain Criteria.

## 4. Discussion

Investigations of aperiodic activity (exponent and offset) are increasingly reported across neurodevelopmental, cognitive, and psychiatric datasets as a putative measure of brain E/I ratios (Molina et al., 2020; Ostlund et al., 2021; Robertson et al., 2019; Thuwal et al., 2021; Wilkinson & Nelson, 2021), with a larger exponent (steeper aperiodic slope) typically interpreted as reflecting increased inhibitory tone (i.e., reduced E/I). This study was the first, to our knowledge, to investigate the extent to which resting state aperiodic activity predicts RDoC *social processes* outcomes in non-clinical young adults.

Given a shift in E/I towards increased excitation has been associated with social communication difficulties in clinical, non-clinical, and animal studies (Bredewold et al., 2015; Cochran et al., 2015; Ford, Nibbs, et al., 2017b; Han et al., 2014), it was predicted that reduced aperiodic exponent (i.e., flatter slope) would be associated with more social communication and processing difficulties. There was a moderate correlation between SRS SCI scores and the aperiodic exponent averaged across all electrodes (i.e., whole brain) indicating that, contrary to expectations, increased inhibitory tone was associated with more social communication difficulties. However, this association did not survive correction for FDR. Furthermore, whole-brain aperiodic exponent did not significantly predict *social processes* outcomes in either the eyes open or eyes closed condition. This may indicate that: 1) predictive power of exponent and offset is possibly region-specific, thus including all electrodes introduces substantial noise that effectively drowns out any signal; 2) there was insufficient power to detect an effect; and/or 3) E/I measured by aperiodic brain activity does not predict the RDoC *social processes* measures utilised here.

To explore the regional specificity of aperiodic activity to *social processes*, two temporo-parietal ‘social brain’ clusters were created for left and right hemisphere that localised to superior, middle, and inferior temporal gyrus, supramarginal gyrus, and angular gyrus (Scrivener & Reader, 2022). These regions are known to be involved in social information processing (Blakemore, 2008; Jung et al., 2019), particularly in the right cortical hemisphere (Abu-Akel et al., 2017; Ford, Nibbs, et al., 2017a; Lindell, 2006; von dem Hagen et al., 2011). Pearson correlations indicated that increased left and right social brain exponent values (more inhibitory tone) were associated with higher SRS SCI scores in our non-clinical sample with a moderate effect size, however, when controlling for false discovery rate, this relationship was no longer statistically significant. There were no associations with RME accuracy or ER-40 performance. Results of the regression models suggested that aperiodic activity within these left and right social brain regions did not significantly predict any of the *social processes* outcomes. Although non-significant, aperiodic exponent in social brain regions were better at predicting SRS SCI scores with moderate effect sizes compared to small effect sizes for whole-brain analysis.

Although there were no significant findings, the trend toward increased exponent (suggestive of greater inhibitory tone) and offset with more *social processes* difficulties warrants further investigation in a larger, more diverse samples, including clinical populations where social communication difficulties are a core feature. Only two studies to date have utilised aperiodic activity to investigate the relationship between brain E/I and social communication – both of which are in clinical young children. Therefore, this study is the first to specifically probe social communication difficulties and E/I via aperiodic activity. Extensive research suggests that social communication difficulties in non-clinical and clinical groups are associated with an increase in E/I (Rubenstein & Merzenich, 2003; Yizhar et al., 2011), implying increased excitation and/or reduced inhibition. However, more recent research has suggested that the nature of the E/I imbalance may be region specific (Ajram et al., 2019; Ford & Crewther, 2016; Nelson & Valakh, 2015), and regional specificity of increased inhibitory tone has been reported in mouse models (J. Gonçalves et al., 2017). The slightly increased aperiodic exponent, potentially reflecting a non-invasive index of greater inhibitory tone in social brain regions, in conjunction with more social communication difficulties observed in this study may reflect underlying neurobiological mechanisms. This is in line with a study in children demonstrating increased exponent with more pronounced autistic traits (Hill et al., 2025) and increased exponent for autistic children with IQ>85 compared to typically developing children and autistic children with IQ<85 (Manyukhina et al., 2022). These findings together indicate that the aperiodic exponent in relation to social communication difficulties should be further examined in larger samples to better understand the nature of neurobiological mechanisms across the autism spectrum.

In line with previous literature, males had a significantly higher aperiodic offset and larger exponent than females in both eyes open and eyes closed conditions (Hill, Clark, et al., 2022; McSweeney et al., 2021). Sex differences in a prior adolescent sample were thought to reflect an advanced neurophysiological maturity in females (Lenroot et al., 2007), which may be the case in this young adult sample as neurodevelopment continues into early adulthood for both males and females (Johnson et al., 2009). Contrary to previous reports (Cellier et al., 2021; Donoghue et al., 2020; Hill, Clark, et al., 2022; McSweeney et al., 2021), exponent and offset were not associated with age, which may reflect that aperiodic activity is relatively stable between 18 and 25 years, or that the age range of the sample was too narrow to detect age effects. Also in line with previous research in children and adolescents, larger exponent and higher offset were recorded in the eyes closed compared to eyes open condition (Hill, Clark, et al., 2022; McSweeney et al., 2021). This reflects a reduction in E/I when eyes are closed, which could be explained by the lack of visual stimulation, attention, and arousal, compared to when participants’ eyes were open (Barry et al., 2009).

There were no significant relationships between RDoC *social processes* self-report (SRS SCI) and behavioural (RME, ER-40) outcome measures themselves. This was somewhat unexpected and possibly problematic given the three measures, SRS SCI, RME, and ER-40, are proposed by the RDoC as gold standard measures of the *social processes* constructs. Although they appear to index different aspects of *social processes*, namely social communication (reception of facial communication [ER-40], and reception and production of non-facial [SRS SCI]), and perception and understanding of others (understanding mental states [RME]), it is reasonable to expect that they would be related. The lack of interrelationship in this study may reflect the limited spread in performance/difficulties in the tasks in this non-clinical sample, insufficient data, or that the measures have poor construct validity, potentially due to poor insight when completing self-report measures. In fact, a recent study demonstrated the RME may lack the psychometric properties to accurately measure theory of mind, compared to the original validation study, and therefore, the lack of associations identified in this study might be due to inadequate measurement tools (Higgins et al., 2023).

The main limitations for this study were the modest sample size (n = 37) and narrow spread of *social processes* difficulties, particularly for SRS scores (range = 42-69). These issues combined may have been a significant contributor to the lack of statistically significant findings, given the small effect sizes reported herein. A more diverse spread of abilities within the sample, for example including clinical participants, may reveal more compelling relationships between aperiodic activity and social processing abilities. Furthermore, EEG is limited in its ability to examine social brain areas specifically due to its relatively poor spatial resolution. Source localization using magnetoencephalography (MEG) and concurrent participant structural brain scans would allow for a more accurate analysis of social brain and *social processes* abilities.

Despite these limitations, these preliminary data show promise for supporting the utility of aperiodic activity in investigating the relationship between brain E/I and social communication difficulties in non-clinical, as well as clinical, populations. Future work should seek to examine this relationship in a larger sample across clinical and non-clinical groups. Furthermore, studies involving pharmacological interventions that modulate excitatory and/or inhibitory processes would provide valuable insights into the utility of the aperiodic exponent as a putative neurobiomarker of *social processes* abilities.

## Acknowledgements

Thank you to Dr Kaila Bianco, Dr Pam Barhoun, and Mr Ji-Shen Loong for assistance in collecting data for this study.

## Data availability

The data that support the findings of this study are available from the corresponding author upon reasonable request.

