## Supplementary Material for "The association between resting state aperiodic activity and Research Domain Criteria Social Processes in young neurotypical adults"

Supplementary Table 1: Descriptive statistics of FOOOF Goodness of Fit

| Metric | N | Mean | SD | Min | Max |
| --- | --- | --- | --- | --- | --- |
| Eyes open R^2^ | 35^a^ | 0.99 | 0.0062 | 0.96 | 1 |
| Eyes closed R^2^ | 37 | 0.99 | 0.0048 | 0.97 | 0.99 |
| Eyes open MAE | 35^a^ | 0.047 | 0.013 | 0.023 | 0.078 |
| Eyes closed MAE | 37 | 0.055 | 0.016 | 0.031 | 0.092 |

Note: R^2^: explained variance, MAE = mean absolute error. ^a^Two participants excluded due to excessive line noise.

**Multivariate Linear Models**

**Whole brain aperiodic activity**

*Outcomes:* Social Responsiveness Scale Social Communication Index (SRS SCI), Penn Emotion Recognition (ER-40) performance, Reading the Mind in the Eyes (RME) accuracy

*Covariates*: Sex

*Model 1 Predictor*: Eyes open exponent

*Model 2 Predictor*: Eyes closed exponent

*Model 3 Predictor*: Eyes open offset

*Model 4 Predictor*: Eyes closed offset

**Social brain aperiodic activity**

*Outcomes*: SRS SCI, ER-40 performance, RME accuracy

*Covariates*: Sex

*Model 1 Predictor*: Left eyes open exponent

*Model 2 Predictor*: Right eyes open exponent

*Model 3 Predictor*: Left eyes open offset

*Model 4 Predictor*: Right eyes open offset
